# The RNA-binding activity of SACSIN HEPN domain is connected to ARSACS

**DOI:** 10.64898/2025.12.18.694329

**Authors:** L. Longo, I. Mariani, A. Cusimano, R. Carrotta, E. Zacco, R. Vazzana, G. Cappelli, M. R. Mangione, M. A. Costa, N. Cuscino, R. Passantino, S. Vilasi, A. Gallo, V. Martorana, R. Giambruno

## Abstract

Autosomal Recessive Spastic Ataxia of Charlevoix–Saguenay (ARSACS) is a neurodegenerative disorder caused by mutations in the *SACS* gene, though the molecular function of its protein product, SACSIN, remains elusive. Therapeutic strategies for ARSACS are limited, mostly due to the exceptionally large size of SACSIN (∼520 kDa), which precludes conventional gene therapy and standard molecular delivery methods.

Over 200 pathogenic variants, including missense and nonsense mutations, have been identified. Regardless of variant type, patients uniformly exhibit cerebellar ataxia and spasticity. Among them, patients carrying missense mutations within the SACSIN higher eukaryotes and prokaryotes nucleotide-binding (HEPN) domain, suggesting that the alteration of the functional role of SACSIN HEPN domain could be linked to the manifestation of ARSACS symptoms. Here, we provide the first direct evidence that SACSIN HEPN domain binds RNA, corroborating the hypothesis that SACSIN functions as an RNA-binding protein. *In silico* predictions through the *cat*RAPID algorithm as well as AlphaFold 3 revealed the propensity of SACSIN HEPN domain to specifically interact with RNAs. These interactions were validated both *in vitro* and in cell studies, confirming that SACSIN HEPN domain binds to RNA transcripts. Overall, our study unveils an unknown molecular function of SACSIN that, if altered, can contribute to the development of ARSACS disease. Therefore, paving the way for innovative therapeutic approaches for ARSACS patients.

## Introduction

Autosomal Recessive Spastic Ataxia of Charlevoix–Saguenay (ARSACS) is caused by mutations in the *SACS* gene^1,2^, which encodes for the multidomain protein SACSIN. Starting from the protein amino terminus, SACSIN possess one ubiquitin-like domain (UBL); three SACSIN internal repeats (SIRPT1, SIRPT1 and SIRPT3) containing three functional subrepeat regions (SR1, SR2 and SR3); one J-domain (DNAJ) and one higher eukaryotes and prokaryotes nucleotide-binding (HEPN) domain^3,4^. The presence of these domains have been associated to the putative co-chaperone function of SACSIN in the cellular protein quality control^5^. Nevertheless, the molecular functions of SACSIN remains still elusive, due to its exceptionally large size that is incompatible with standard biochemical assays. Cells and mice depleted or with a reduced expression of SACSIN show aberrant alterations of the mitochondrial functions, improper microtubules and microfilaments formation, and accumulation of protein aggregates with the consequent induction of autophagy^6^. It is still under debate whether these alterations are cause or consequence of the absence of SACSIN biological functions, especially as these altered processes are common to many neurodegenerative disorders^7^.

So far, more than 200 mutations have been identified in ARSACS patients along the whole *SACS* gene sequence, including missense and stop codon mutations. Irrespective of the type of mutation, these patients show the canonical pathophysiological phenotypes of ataxia and spasticity^4^. Among them, ARSACS patients carrying missense mutations in the SACSIN HEPN domain (N4463K, Y4490S, E4511NTer9, N4549D, F4574C), which is exactly at the C-terminus of the protein^8–12^; thus, suggesting that the manifestation of ARSACS symptoms may be connected to the absence or the altered functions of the HEPN domain.

The HEPN domain, present both in prokaryotes and eukaryotes, was initially reported to bind to nucleotides and serve as substrate-binding subunit of minimal nucleotide-transferases (MNT)^13^. Afterwards, all proteins containing a HEPN domain were grouped into the HEPN family, which is a class of RNA-binding proteins (RBPs). Most of them, having an endoribonuclease activity, belong to a subset named HEPN RNase^14^. HEPN RNases contain a conserved RΦxxxH motif within their HEPN domain that constitutes a metal-independent endoRNase active site^14,15^. Other HEPN family proteins, such as the human SACSIN, lacking the RΦxxxH motif have been suggested to function as non-catalytic RBPs^15^.

The concept that SACSIN may potentially bind to cellular RNA transcripts through its HEPN domain led us to hypothesize that alterations in SACSIN RNA-binding properties could be linked to several molecular defects observed in ARSACS patient cells.

Taking advantage of *in silico, in vitro* and in cells experiments, we are proving for the first time the ability of SACSIN HEPN domain to bind to RNA transcripts. Specifically, we discovered that HEPN binds to both coding and non-coding RNA transcripts. SACSIN HEPN RNA-binding activity seems to have a specific role in the proximity of the endoplasmic reticulum (ER), where it interacts with the 7SL RNAs composing the signal recognition particle (SRP) complex (*e. g.* RN7SL1). The SRP is a ribonucleoprotein complex recognizing and targeting new proteins to the ER for their subsequent cellular trafficking^16^. Overall, suggesting a new functional role for SACSIN that could explain the protein trafficking dysfunction observed in ARSACS cells^17^.

Moreover, the interaction between SACSIN HEPN domain and the RNA is necessary to compartmentalize the protein in close proximity to the ER and to regulate its propensity to form protein condensates. Protein condensates are reversible membraneless assemblies of proteins forming in cells through liquid-liquid phase separation process (LLPS)^18,19^. The RNA can either act as a scaffold promoting LLPS or as negative regulator for the formation of protein condensates^20–22^. Here, we found out that SACSIN HEPN domain forms protein condensates when its RNA substrates are not available.

## Results

### SACSIN HEPN domain binds to RNA

The ability of SACSIN to bind to RNA has never been tested, although SACSIN HEPN domain can bind to guanosine triphosphate (GTP) with low micromolar affinity^23,24^.

We initially investigated the literature and found out that two independent studies, relying on the organic-aqueous phase separation methodology to unbiasedly purify and detect RBPs from cells, identified SACSIN among the proteins potentially able to interact with RNA transcripts in human cells^25,26^. Consistently, we observed that SACSIN formed high molecular weight complexes upon UV-crosslinking (**Supplementary Figure 1A**), as reported for other RBPs^27^; hence, confirming the ability of SACSIN to interact with cellular RNAs.

We investigated SACSIN propensity to interact with cellular RNA transcripts through the well-established protein-RNA interaction prediction algorithm *cat*RAPID (**Figure 1**)^28^. Interestingly, both the UBL and the HEPN domain, respectively located at the amino- and carboxy-terminus of SACSIN, were predicted to have the ability to bind to RNA, with an overall interaction propensity ≥ 0.6 (**Figure 1A**). Neither domain showed a preference for a specific RNA-binding motif (**Figure 1A**). For the UBL domain, no discrete amino acid region exceeded the RNA-binding threshold of 0.5. In contrast, the analysis of the HEPN domain identified a 50-amino acid region (V4474-V4524) as the main contributor to RNA binding, with the highest predicted interaction propensity > 0.65 (**Figure 1B**).

**Figure 1.**
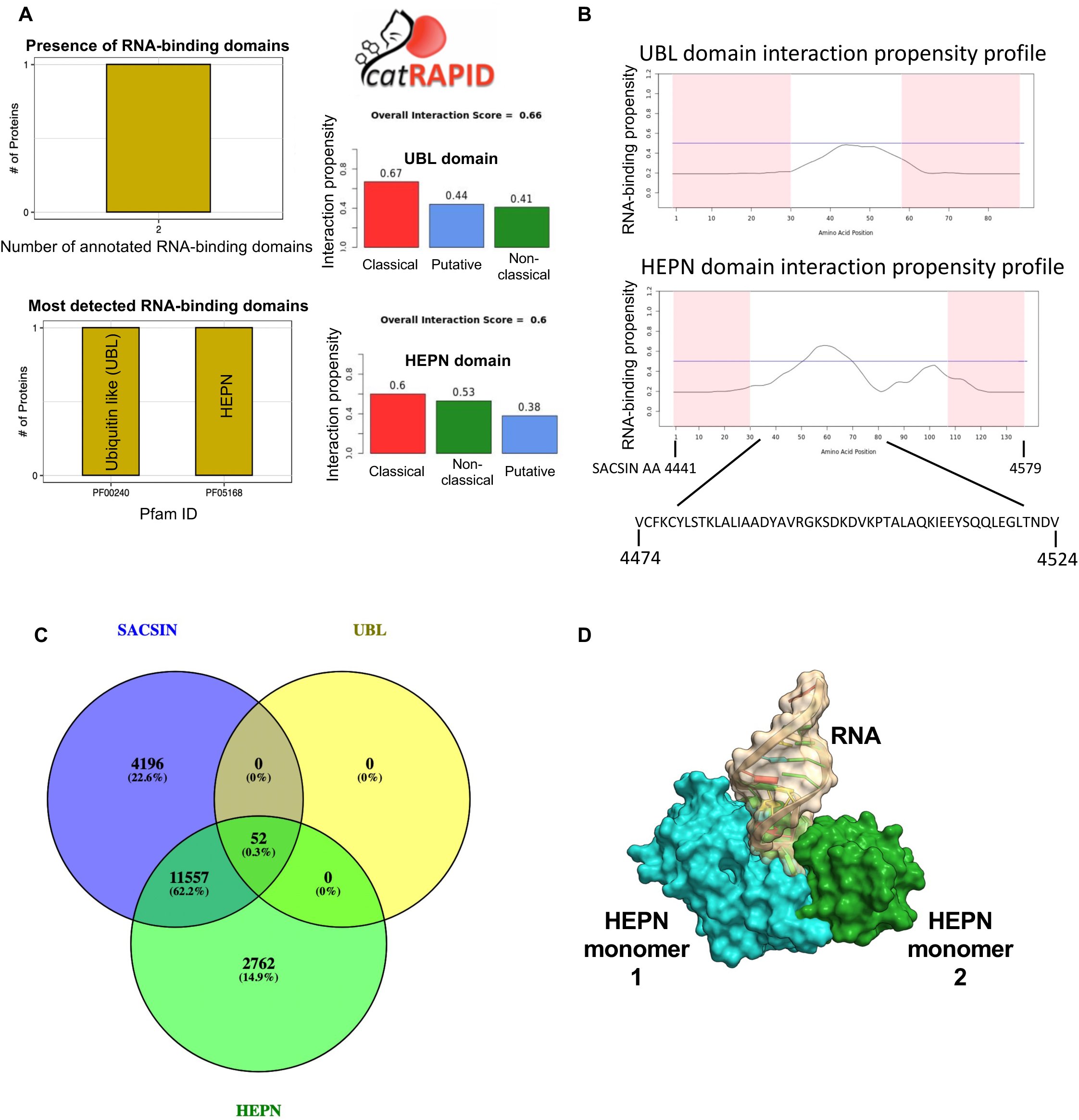
In silico predictions of SACSIN HEPN - RNA interactions. **A)** Left. *cat*RAPID prediction of the interaction propensity of SACSIN toward cellular RNA transcripts. Two SACSIN domains (UBL and HEPN) are prone to bind to RNA. Right. Interaction propensity scores calculated by the *cat*RAPID algorithm for the UBL (upper part) and HEPN (lower part) domain. A positive interaction propensity is considered when Z-score > 0.5. **B)** Interaction propensity profile of SACSIN UBL (upper part) and HEPN (lower part) domain calculated trough the *cat*RAPID algorithm. For the UBL domain, no specific binding regions were detected within the domain. The HEPN domain, instead, harboured a region of 50 amino acids (SACSIN amino acids (AA) 4474- 4524) with high interaction propensity with RNA transcripts. **C)** Venn diagram of the human protein-coding RNAs predicted by *cat*RAPID as putative interactors of either SACSIN, the UBL or the HEPN domain. Only protein-coding RNAs with a Z-score > 0.5 were considered. **D)** AlphaFold 3 prediction of the HEPN domain in its dimeric form interacting with a predicted GC-rich RNA sequence.

We then used *cat*RAPID to predict the human RNA transcripts interacting with the full-length SACSIN, as well as its two putative RNA-binding domains, UBL and HEPN. Positive RNA-protein interactions were selected by applying a dynamic *cat*RAPID Z-score of > 0.5, as previously reported^28^. Full-length SACSIN and the HEPN domain showed the highest number of predicted protein-RNA interaction pairs, while few RNAs were expected to associate with the UBL domain (**Figure 1C**). Overall, suggesting that HEPN is the domain of SACSIN required for the interaction with RNAs. More than 60% of the putative interacting protein-coding RNAs were shared between full-length SACSIN and the HEPN domain (**Figure 1C**), while the overlap among non-coding RNAs was lower (**Supplementary Figure 1B**). We therefore focused on protein-coding RNAs with high interaction propensity (Z-score > 3.5) for both SACSIN and the HEPN domain and performed sequence alignment analysis using Clustal Omega Multiple Sequence Alignment algorithm^29^. Interestingly, we found the presence of a specific GC-rich consensus sequence, which was absent in the non-interacting RNAs (*cat*RAPID Z-score < 0) (**Supplementary File 1**). We analyzed the interaction of the predicted RNAs with the dimeric form of the HEPN domain, as we observed that the recombinant protein form stable dimers (**Supplementary Figure 1C**), in line with the literature^23^. We used AlphaFold 3^30^ to predict the structure of complexes between the HEPN dimer and a series of RNA sequences that were found by *cat*RAPID (**Supplementary File 1**). Notwithstanding the well-known AlphaFold 3 limitations in reliably predicting RNA structure, we were able to single out an interesting complex structure interaction with a GC-rich RNA sequence, indicating a specific interaction with the dimer (**Figure 1D, Supplementary Figure 1D and E)**. We confirmed the stability and specificity of the HEPN-RNA interactions by molecular dynamics simulation, testing the stability of the complex along a 500 ns trajectory. Analogous molecular dynamics simulation with a GC-poor RNA sequence of the same length showed, instead, a weak and non-specific interaction with the HEPN dimer. In this latter case, the RNA molecule was displaced from the initial position and visited various regions of the dimer surface (**Supplementary Figure 1D and E**). The interacting RNA localized within a hydrophilic cleft present at the monomer-monomer interface of the HEPN dimer (**Figure 1D**), mirroring the interaction binding mode of SACSIN HEPN with GTP, previously observed by NMR spectroscopy^23^.

We validated the interaction *in vitro* through GST-pulldown assays with recombinant GST-tagged SACSIN HEPN and UBL-SR1 proteins^23^ and a synthetic biotinylated GC-rich RNA oligo of 32 nucleotides retrieved from *cat*RAPID predictions (**Supplementary Figure 1E and Supplementary File 1**). The biotinylated RNA oligo was efficiently pulled down by GST-HEPN, while no interaction was detected with the GST-UBL-SR1 (**Figure 2A**), confirming *cat*RAPID algorithm predictions. We then produced the recombinant HEPN domain devoid of GST-tag and measured its affinity for the RNA oligo using biolayer interferometry (BLI). The HEPN domain bound the predicted RNA target with a dissociation constant (K_D_) of 2.6 ± 0.3 µM (**Figure 2B**), demonstrating that this domain recognizes the RNA with measurable affinity. No significant changes in the secondary structure of either the protein or the RNA were observed by circular dichroism (CD) (**Supplementary Figure 2A and B**). Likewise, no variations were observed in the protein tertiary structure monitored by the intrinsic fluorescence emission of HEPN in the presence of RNA (**Supplementary Figure 2C)**. Overall, indicating that no structural perturbation occurs upon binding.

**Figure 2.**
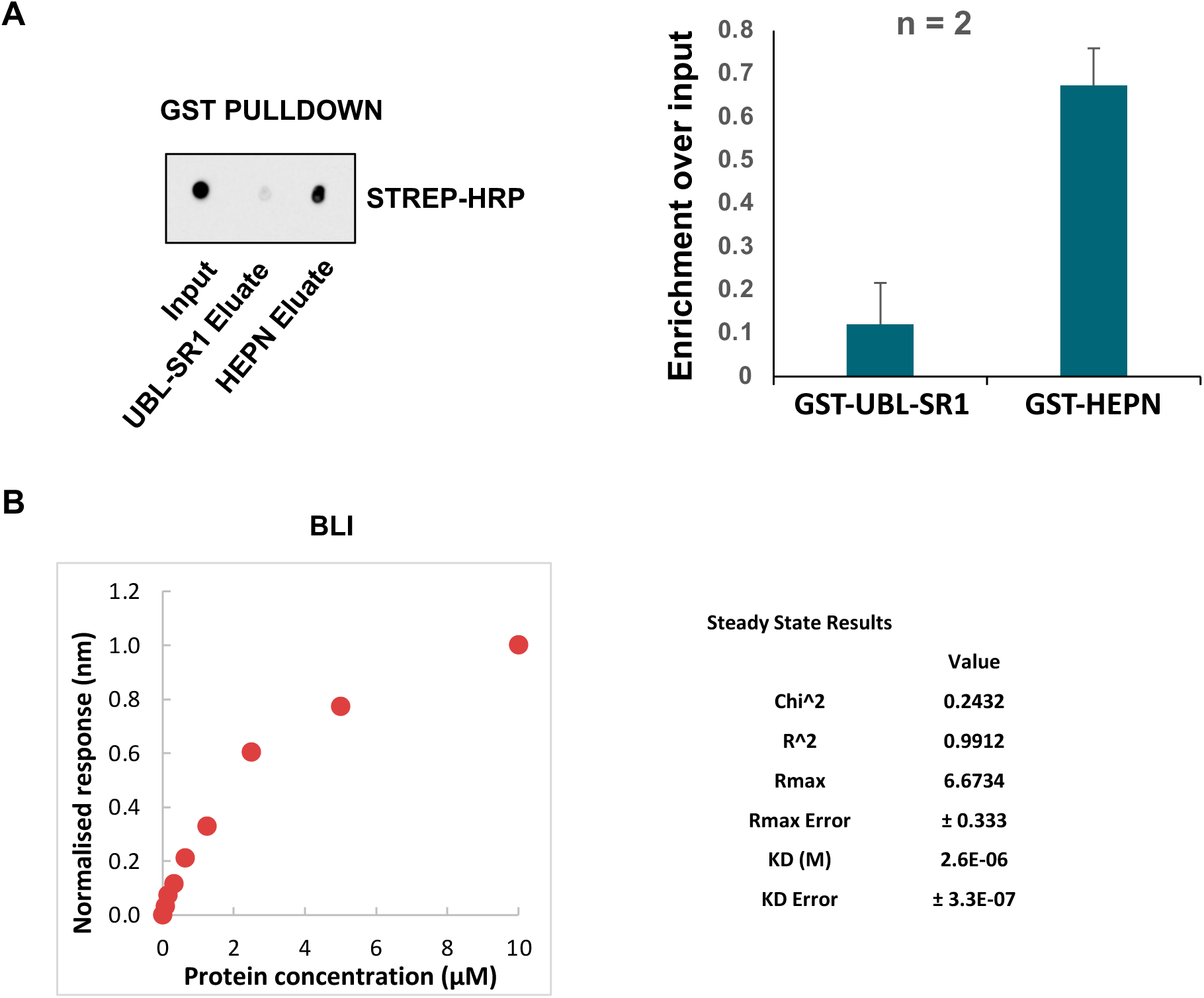
In vitro experiments to demonstrate SACSIN HEPN RNA-binding ability. **A)** GST-pulldown of recombinant GST-tagged proteins with the synthetic biotinylated GC-rich RNA sequence. The data showed the average and the standard deviation of two biological replicates (n = 2). **B)** BLI experiments to assess the K_D_ of GST-HEPN with the synthetic biotinylated GC-rich RNA sequence. Data were collected from three independent experiments and the average results are reported.The results were normalized by setting the highest value of the binding response to 1.

### SACSIN HEPN domain localizes to the ER, where it interacts with the SRP complex

We sought to identify the cellular RNA transcripts bound by the HEPN domain in human cells using the APEX-Seq strategy^31,32^. This proximity biotinylating system enables the identification of the RNA interactors of a specific protein in living cells, without the use of crosslinking agents^33^. We fused FLAG-APEX2 to the N-terminus of the HEPN domain as well as to a control eGFP, generating two fusion proteins of 46 and 60 kDa, respectively. The APEX2-fusion proteins were expressed from a tetracycline-inducible mammalian expression vector integrated in a single active genomic locus of HeLa Flp-In^TM^ TRex cells; hence, permitting a homogenous and basal expression level of the exogenous FLAG-APEX2-tagged proteins. Upon doxycycline administration, the expression of FLAG-APEX2-GFP and FLAG-APEX2-HEPN was comparable and remained stable over time (**Figure 3A**). In line with the expression of the endogenous SACSIN, FLAG-APEX2-HEPN localized to the cytoplasm. Interestingly, we observed that the protein was also present in the proximity of the endoplasmic reticulum (ER), especially after 72h of doxycycline administration (**Figure 3B and Supplementary Figure 3**). In contrast, FLAG-APEX2-GFP remained widely distributed within the cellular compartments (**Supplementary Figure 3**). FLAG-APEX2-GFP was promiscuously associated with RNA transcripts, as previously reported^32^. FLAG-APEX2-HEPN, instead, began to interact with cellular RNA transcripts only after 48h of doxycycline administration (**Figure 3C**). No biotinylated RNA signal was detected after 24h of doxycycline administration, as well as in the absence of biotin (**Figure 3C**). To identify the RNA transcripts bound by FLAG-APEX2-HEPN, we performed APEX-Seq in biological triplicates in cells expressing either FLAG-APEX2-HEPN or FLAG-APEX2-GFP, after 72h of doxycycline administration (**Supplementary Figure 4A**). The resulting biotinylated RNAs were affinity purified using streptavidin beads (**Supplementary Figure 4B**) and sequenced through the Illumina next generation sequencing platform. The expression of FLAG-APEX2-HEPN did not alter the cellular transcriptome (**Supplementary Figure 5A**), with only 8 de-regulated RNA transcripts detected (**Supplementary File 2**). A total of 67 RNAs specifically and reproducibly interacted with FLAG-APEX2-HEPN compared to FLAG-APEX2-GFP (**Figure 3D and Supplementary File 2**). These data were intersected with *cat*RAPID predictions to retrieve the *bona fide* list of HEPN RNA interactors. Among the interacting RNA transcripts, we found that 31 RNAs have a positive *cat*RAPID score (Z-score > 1.2) (**Figure 3E**). The remaining 36 RNA transcripts were either absent from the *cat*RAPID library (26 RNA transcripts) or predicted not to interact with HEPN or SACSIN (Z-score < 1; 10 RNA transcripts). The non-predicted RNA interactors included Ces1 and S100p that were transcriptionally upregulated in FLAG-APEX2-HEPN cells (**Supplementary File 2**). For this reason, we considered the non-predicted RNA interactors as false positives. The final list of *bona fide* HEPN interactors was composed of 57 RNA transcripts, comprising 28 mRNAs, 27 non-coding RNAs and 2 mitochondrial RNAs (**Supplementary File 2**). Gene ontology enrichment analysis revealed a significant enrichment of ER-localized RNAs and RNAs composing the SRP ribonucleoprotein complex, which regulates cellular protein trafficking^16^ (**Supplementary File 2**). These results suggest that the HEPN domain participates in cellular protein trafficking, a process severely impaired in ARSACS cells^17^. Finally, by ultraviolet-RNA immunoprecipitation (UV-RIP-FLAG), we confirmed that FLAG-APEX2-HEPN specifically interacted with the RNA transcript coding for RPL26 and the non-coding RNA RN7SL1, which is the main core RNA component of the SRP complex^16^ (**Figure 3F**).

**Figure 3.**
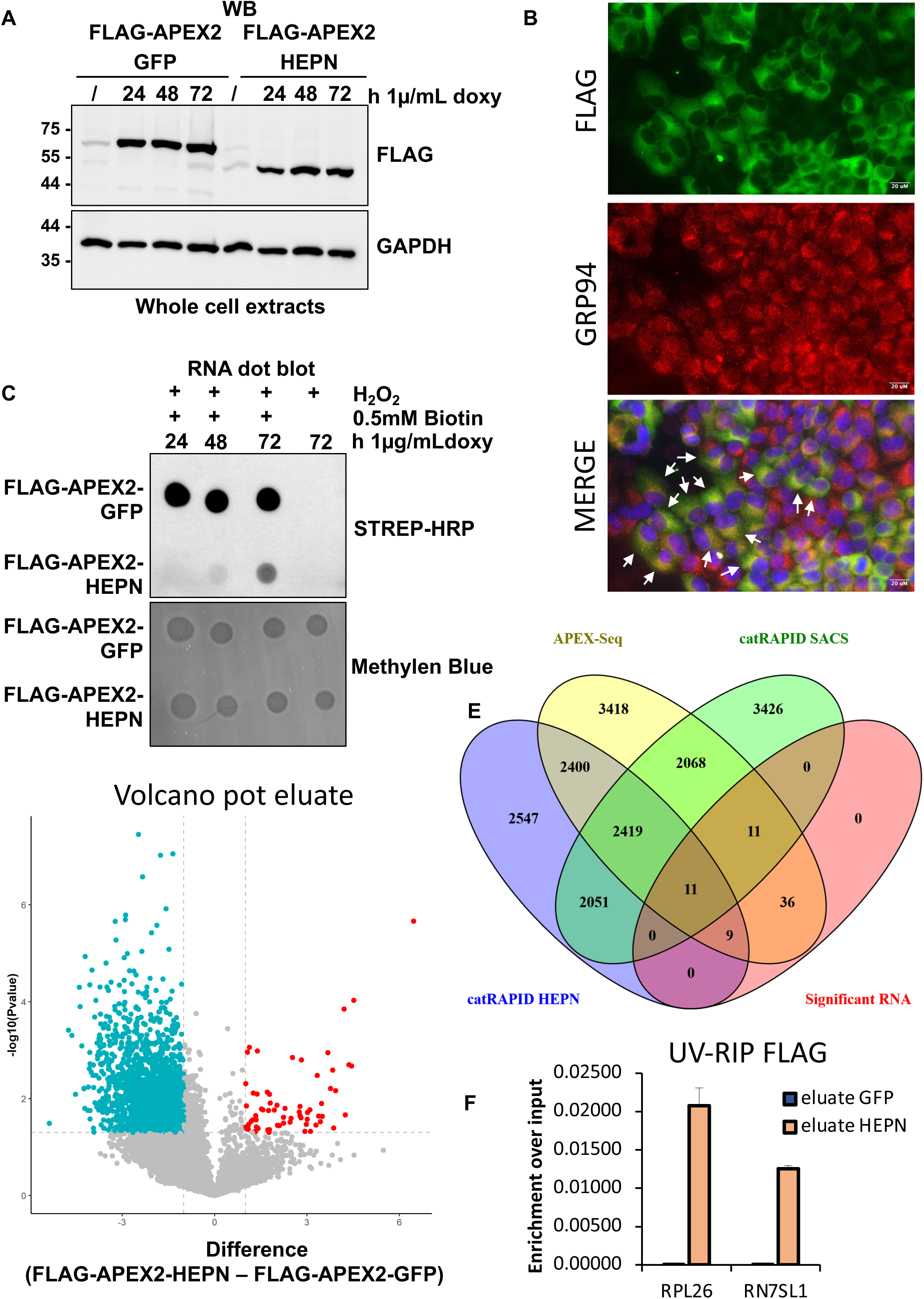
Confirmation of SACSIN HEPN RNA-binding activity and identification of RNA targets. **A)** Expression of FLAG-APEX2-GFP and FLAG-APEX2-HEPN over time in response to doxycycline administration. GAPDH was used as loading control. The displayed figure is the representative image of one of the three performed biological replicates (n = 3). **B)** Immunofluorescence analysis of Hela Flp-In^TM^ T-Rex cells expressing FLAG-APEX2-HEPN, upon 72h 1µg/mL doxycycline administration. In green, the signal for FLAG-APEX2-HEPN while in red the protein GRP94, which is a chaperone protein located in the endoplasmic reticulum (ER). The white arrows indicated the cells where we observed a co-localization of the two proteins, confirming that FLAG-APEX2-HEPN accumulated in the ER-proximity. The displayed figure is the representative image of one of the two performed biological replicates (n = 2). **C)** RNA dot blot analysis of the biotinylated RNAs recovered in response to the biotinylation reaction for the FLAG-APEX2-GFP and FLAG-APEX2-HEPN proteins at different time of doxycycline administration. Methylen blue staining was used as loading control of the RNAs spotted into the nitrocellulose membrane. The displayed figure is the representative image of one of the three performed biological replicates (n = 3). **D)** Volcano plot analysis of streptavidin pulldown eluate derived from FLAG-APEX2-HEPN and FLAG-APEX2-GFP cells. RNA transcripts significantly enriched (log_2_ Fold change (Fc) > 1; two-sided T-Test Pvalue < 0.05) in FLAG-APEX2-HEPN and FLAG-APEX2-GFP cells are colored in red and blue, respectively. **E)** Venn diagram analysis of i) the significant RNA transcripts associated with FLAG-APEX2-HEPN compared to FLAG-APEX2-GFP identified by APEX-Seq (Significant RNA; red ellipse); ii) all the RNA transcripts found associated with FLAG-APEX2-HEPN by APEX-Seq (APEX-Seq; yellow ellipse); iii) RNA transcripts predicted to associate with SACSIN (*cat*RAPID SACSIN; green ellipse); iv) RNA transcripts predicted to associate with SACSIN HEPN domain (*cat*RAPID HEPN; violet ellipse). **F)** UV-RNA Immunoprecipitation (UV-RIP)-FLAG of HeLa Flp-In^TM^ T-REx cells expressing either FLAG-APEX2-GFP or –HEPN, in response to 72h of 1µg/mL doxycycline administration. We confirmed that the RNA transcript coding for RPLP26 and the non-coding RNA RN7SL1 were interacting with the immunopurified FLAG-APEX2-HEPN, in agreement with the APEX-Seq data. The data showed the average and the standard deviation of three technical replicates.

### The interaction with the RNA inhibits the formation of SACSIN HEPN condensates

Since SACSIN HEPN domain binds to RNA, we sought to investigate whether it has a propensity to form RNA-dependent protein condensates^20–22^. Hence, we performed *in vitro* phase separation assays using the crowding agent polyethylene glycol PEG6000^34,35^. Light scattering at 30° revealed a time-dependent growth of HEPN condensates (**Figure 4A and 4B)**. Thioflavin-T staining confirmed that the final aggregates lacked amyloid-like properties (**Figure 4C)**. Interestingly, static light scattering showed that the presence of total cellular RNA transcripts inhibited protein condensates formation in a dose-dependent manner (**Figure 4A)**. Furthermore, Multi Angle Light Scattering (MALS) analysis revealed RNA-dependent structural differences in the form factor of the protein condensates formed by HEPN alone compared to the ones formed by HEPN in the presence of RNA. The data recorded for HEPN alone were best defined by the presence of large spherical species with a diameter of ∼2.6 μm, consistently with optical microscopy inspection of the sample where protein assemblies of microns size were detected **(Figure 4A and 4B)**. Whereas in the presence of RNA, much smaller spherical species with a maximum diameter of 360 nm were detected (**Figure 4A)**.

**Figure 4.**
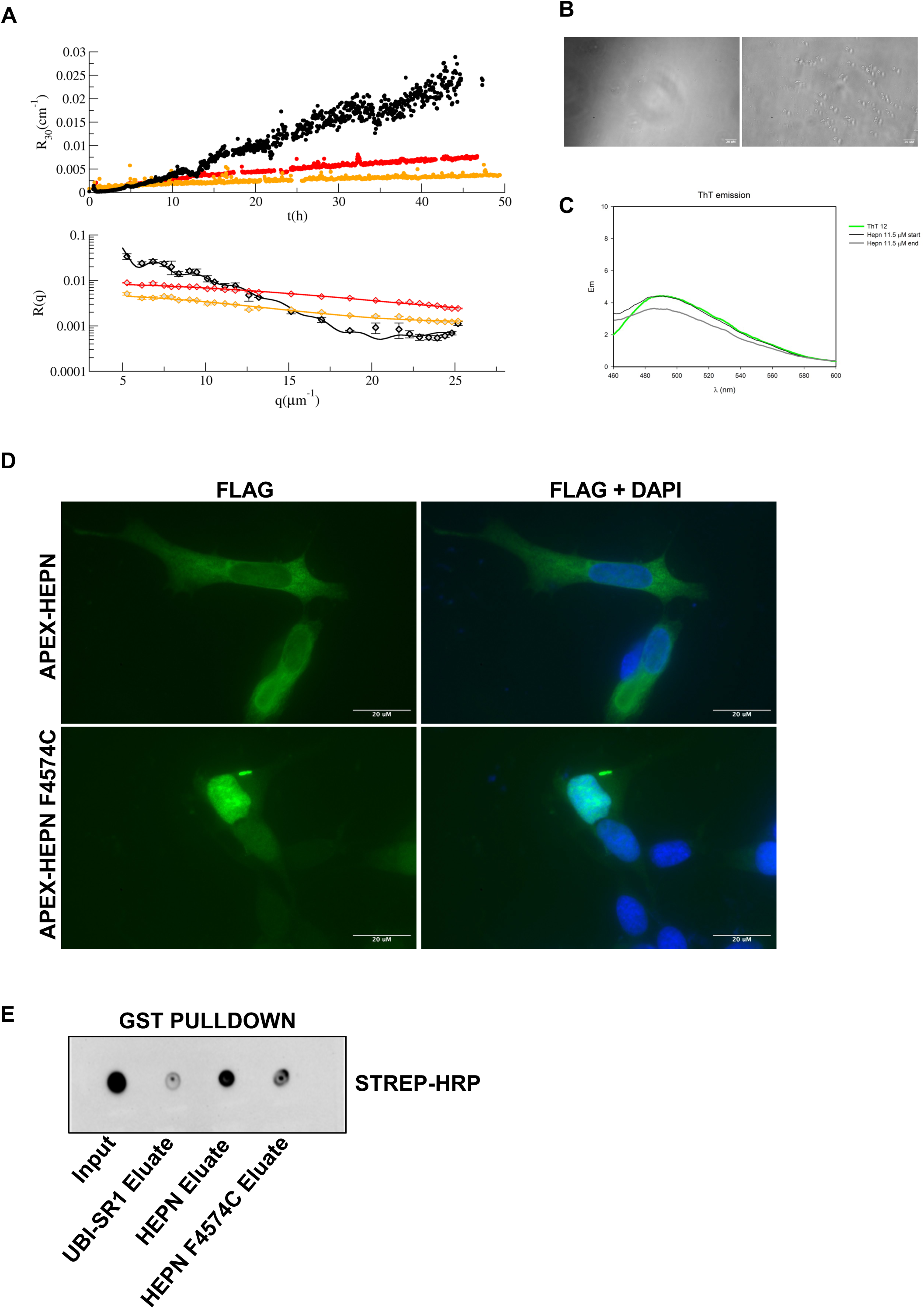
SACSIN HEPN RNA-binding properties regulates its propensity to form protein condensates. **A)** Time-lapse Rayleigh ratio at θ=30° of SACSIN HEPN domain alone (black curve) or mixed with 70 μg (low RNA, red curve) or 77 μg (high RNA, orange curve) of total RNA extracted from human cell lines, in all cases in the presence of 15% PEG6000 (top panel). R (q) MALS data for the three samples at the end of the kinetics, represented with the data best fitting functions (continuous lines) obtained according to the model reported in the material and methods section. HEPN forms larger condensates that did not occur in the presence of the RNA (bottom panel). **B)** Representative bright-field images of SACSIN HEPN domain condensates using a wide-field microscope 40x magnification. **C)** Thioflavin (ThT) assay of SACSIN HEPN domain at the beginning and at the end of the kinetic. No significant variations in the ThT emission signal were observed between the initial and final samples, assessing that the HEPN condensates are not amyloid fibrils. **D)** Immunofluorescence analysis of SACSIN knock-out SH-SY5Y cells transiently expressing FLAG-APEX2-HEPN or FLAG-APEX2-HEPN F4574C and fixed after 24h from transfection with 4% PFA. Subsequently, immunofluorescence staining was performed with a-FLAG antibody. DAPI was used to stain the nuclei. The displayed figure is a representative image of the three performed biological experiments (n = 3). **E)** GST-pulldown of recombinant GST-tagged proteins with the synthetic biotinylated GC-rich RNA sequence. The displayed figure is the representative image of one of the two performed biological replicates (n = 2).

We next established a doxycycline-inducible system for live-cell imaging of GFP-HEPN and observed that the overexpressed protein was initially soluble and localized in the cytoplasm (**Supplementary Figure 5B**). Over time, it translocated to the nucleus, where it formed protein condensates (**Supplementary Figure 5C**). In contrast, GFP-HEPN carrying the ARSACS-associated F4574C mutation was immediately mis-localized to the nucleus and formed protein condensates at early time points (**Supplementary Figure 5B**). Similar results were obtained in SACSIN knock-out SH-SY5Y and HeLa cell lines transiently overexpressing FLAG-APEX2-HEPN or FLAG-APEX2-HEPN F4574C (**Figure 4D, Supplementary Figure 6 and 7A**). Based on these results, we hypothesized that the F4574C mutation might impair the RNA-binding properties of SACSIN. Consistently, GST-HEPN F4574C showed reduced RNA-binding ability compared with the wild-type protein (**Figure 4E**). However, we were unable to assess the RNA-binding properties of SACSIN HEPN F4574C in cells because the protein was sensibly less expressed than SACSIN HEPN (**Supplementary Figure 7B**).

## Discussion

Our study provides the first direct evidence that SACSIN can bind cellular RNAs through its HEPN domain, revealing a previously uncharacterized molecular function that had been computationally predicted^15^ but never experimentally investigated. Using the APEX2-tagged HEPN domain, we reported that SACSIN RNA-binding is functionally linked to the phenotypic defects in protein trafficking observed in ARSACS cells. The HEPN domain interacts with the SRP ribonucleoprotein complex^16^, a critical mediator of ER-targeted protein translocation^36,37^, suggesting that SACSIN may act as a local coordinator of nascent peptide recognition and ER trafficking. These findings offer a mechanistic framework for understanding the impaired trafficking of membrane and secreted proteins observed in ARSACS cells^17^.

We further show that SACSIN HEPN domain exhibits a propensity to form protein condensates, a behaviour modulated by its RNA-binding properties. This observation suggests a dual role for the RNA: i) a structural scaffold promoting proper subcellular localization near the ER; ii) a chaperone function preventing aberrant homodimeric interactions. Disruption of SACSIN HEPN RNA-binding ability, as exemplified by the F4574C mutation, leads to aberrant nuclear condensation, potentially progressing toward toxic protein aggregates. This mechanism echoes pathogenic processes observed in other neurodegenerative disorders, including amyotrophic lateral sclerosis (ALS) and frontotemporal lobar degeneration (FTLD), where mutations in RNA-binding proteins such as FUS and TDP-43 drive pathological condensation and aggregation^38–40^.

Our data also raise important considerations regarding genotype-phenotype correlations in ARSACS patients. While most patients exhibit near-complete loss of SACSIN due to co-translational ubiquitination^41^, individuals harbouring C-terminal mutations express truncated proteins^12^ that may accumulate as condensates, potentially forming toxic aggregates. This suggests that distinct molecular mechanisms may underlie similar clinical phenotypes, highlighting the potential diagnostic value of detecting SACSIN protein aggregates in differentiating ARSACS subtypes.

Critically, although our findings establish SACSIN as an RNA-binding protein with potential roles in protein trafficking and condensate regulation, several questions remain. The precise RNA targets and the structural determinants mediating SRP interaction require further characterization. Moreover, the long-term cellular consequences of SACSIN condensates, including their possible transition to irreversible aggregates, warrant in-depth investigation. Future studies employing ARSACS cellular and animal models, particularly using the compact HEPN domain, may pave the way for targeted therapeutic strategies aimed at restoring protein trafficking or preventing aggregate formation.

In conclusion, our work not only uncovers a previously unappreciated molecular function for SACSIN but also links RNA-binding activity to disease-relevant cellular phenotypes, providing a conceptual framework for understanding ARSACS pathogenesis and offering potential avenues for therapeutic intervention.

## LIMITATION OF THE STUDY

Our findings are mainly addressing the molecular function of SACSIN HEPN domain, while we could not collect the same experimental data with the full-length protein due to experimental limitations. Working with the endogenous SACSIN is extremely challenging and we will require additional different experimental setup to address its RNA-binding properties. Moreover, some of SACSIN-RNA interactions may be tissue specific and part of our findings will have to be confirmed in neuronal differentiated cells as well as in ARSACS animal models.

## Materials & methods

### Reagents and plasmids

The following plasmids were purchased from Addgene: pDEST-cDNA5-FRT/TO-3*Flag-APEX2 N-term was a gift from Benjamin Blencowe (Addgene plasmid # 182925; http://n2t.net/addgene:182925; RRID: Addgene_182925); pGEX4T-HEPN was a gift from Kalle Gehring (Addgene plasmid # 75296; http://n2t.net/addgene:75296; RRID:Addgene_75296); pGEX6P1-Ubl-sr1(SIRPT1) was a gift from Kalle Gehring (Addgene plasmid # 75298; http://n2t.net/addgene:75298; RRID:Addgene_75298). pRetroX GFP Mps1 wt was a gift from Floris Foijer (Addgene plasmid # 63702; http://n2t.net/addgene:63702; RRID: Addgene_63702)

The cDNA of the HEPN domain was PCR-amplified using the following primers: Forward 5’-GGGGACAAGTTTGTACAAAAAAGCAGGCTCCGTTGGCAATCCAGTGGAAGC-3’;

Reverse 5’-GGGGACCACTTTGTACAAGAAAGCTGGGTTTACACTTTTTGTTGCATAAAATTTT CAAGT- 3’.

The amplified cDNA was cloned into the pDonor201 through BP reaction of the gateway cloning strategy (11789020, ThermoFisher Scientific). The cDNA was then transferred by LR reaction of the gateway cloning strategy (11791020, ThermoFisher Scientific) into the pDEST-cDNA5-FRT/TO-3*Flag-APEX2 N-term.

For the generation of pRetroX GFP-HEPN, both pRetroX GFP Mps1 wt and pGEX4T-HEPN were digest with BamHI (BA3136S, New England Biolabs) and NotI (BA3189S, New England Biolabs). The digested products were run on an agarose gel and the respective linear bands of insert (HEPN) and backbone (pRetroX GFP) were extracted from gel using the Wizard(R) SV Gel and PCR Clean-up System (A9281, Promega). The bands were then ligated overnight using the T4 DNA ligase (M0202S, New England Biolabs).

The HEPN F4574C mutant was generated by Q5® Site-Directed Mutagenesis Kit (BE0552S, New England Biolabs) using the following primers:

Forward 5’-ACTTGAAAATtgtATGCAACAAAAAGTG– 3’; Reverse 5’ - TTTATTATGATACAGGCAGTAC– 3’.

All the plasmids have been sequence verified by the sanger sequencing service offered by Eurofins.

Synthetic RNA oligos of 32 nucleotides were synthesized by Eurofins according to the following sequences: Predicted HEPN interacting RNA: CGGGGGCGGCGGCGGCGGCGGCGGCAGCGGGG.

For the GST pulldown and BLI assays, the predicted HEPN interacting RNA was synthesized including a 5’ biotin.

### Cell lines

HeLa, HEK293T and HeLa Flp-In^TM^ TREx cells were grown in DMEM with glucose and L-glutamine (ECM0103L, Euroclone), 100 U/mL penicillin and streptomycin (ECB3001D, Euroclone) and supplemented with either 10% fetal bovine serum (FBS) (ECS5000L, Euroclone) or tetracycline free FBS (ECS01822, Euroclone), respectively.

SACSIN knock-out SH-SY5Y cells, kindly provided by Prof. F. M. Santorelli, were grown as above described except that cultivation medium was complemented with 15% FBS and supplemented with non-essential amino acids (ECB3054D, Euroclone).

### APEX-Seq

Cells were incubated at 37 °C for 30□minutes with complete media containing 0.5□mM biotinyl-tyramide (SML2135, Merck Life Science). Afterwards, the medium was discarded and 1 mM of H_2_O_2_ (H1009, Merck Life Science) in PBS was provided for 1 minute at room temperature followed immediately by 3 washes with freshly prepared quenching solution (10 mM sodium ascorbate (27688.235, VWR Chemicals), 5 mM Trolox (238813, Merck Life Science), 10 mM sodium azide (S2002, Merck Life Science), PBS 1x). Cells were then pelleted and RNA extracted using the Miny RNAse extraction kit (R1054, Zymo Research), following vendor’s instruction and including the in-column DNAse I digestion step.

The eluted material was analyzed by RNA-dot blot assay using a STREP-HRP conjugate to confirm the presence of biotinylated RNAs. For each sample, 60 μg of total RNA was diluted with 1 mL of Blocking Buffer (5x Denhardt’s Solution (D9905-5ML, Merck Life Science) in PBS1x supplemented with RNAse inhibitor) and 10% of the solution was taken as input material. The remaining 90% was affinity purified rocking it for 1 hour at room temperature in combination with 30 uL of MyOne C1 Streptavidin Dynabeads (65001, ThermoFisher Scientific), that were previously washed 3 times with Buffer 1 (0.5% sodium deoxycholate in PBS 1x supplemented with RNAse inhibitor (BIO-65028, Meridian Bioscience) and equilibrated with Blocking Buffer for 30 minutes at room temperature on a nutator. Samples were then placed on a magnetic rack to separate the unbound fraction from the magnetic beads. Beads were subsequentially washed with four different washing buffers (Buffer 2: 6 M urea pH 8.0, 0.1% SDS in PBS; Buffer 3: 2% (w/v) SDS in PBS; Buffer 4: 750 mM NaCl, 0.5% (w/v) sodium deoxycholate, 0.1% (w/v) SDS, in PBS; Buffer 5: 150 mM NaCl, 0.5% (w/v) sodium deoxycholate, 0.1% (w/v) SDS, in PBS) to maintain only the biotinylated RNA bound to the beads. A total of 2 washes of 1 mL of each buffer was performed. At the end of the washes, the bound biotinylated RNA was extracted from the beads using the Quick-RNA MicroPrep Kit (R1050, Zymo Research) and the RNA was eluted with 14 μL of nuclease free water.

### RNA dot blot assay

RNA was spotted on hybound nitrocellulose membrane (GERPN203B, Merck Life Science) and let it adsorb into the membrane. Then, the RNA was UV-crosslinked to the membrane using the UV-stratalinker with one cycle of 2500 Kjoule. To reveal the presence of biotinylated RNA, the membrane was then stained with the Chemiluminescent Nucleic Acid Module kit (89880, ThermoFisher Scientific), following vendor’s instruction, Afterwards, the membrane is stained for 5 minutes with 0.1% (w/v) Methylene Blue solution in 0.3 M sodium acetate (pH 5.2. The pH is adjusted with acetic acid) and de-stained with milliQ water.

### Library preparation and Next-generation sequencing

RNA integrity was evaluated with the Agilent 4200 TapeStation System (Agilent Technologies Ltd). RNA-Seq libraries for Input samples were prepared starting from 500 ng of RNA using the Illumina TruSeq Stranded Total RNA (20020596, Illumina) and TruSeq RNA UD Indexes 24 Indexes-96 Samples (20020177, Illumina). Briefly, total RNA input was deprived of ribosomal RNA and fragmented before the synthesis of single and double strand cDNA, as previously described^42^. Then, double strand cDNA 3’ends were adenylated and Illumina indexing adapters were ligated. RNA-Seq libraries for HEPN-Elute and GFP-Elute samples were prepared using the NEBNext Ultra II Directional RNA Library Prep Kit for Illumina (BE7760S, New England Biolabs) and NEBNext Multiplex Oligos for Illumina (Index Primers Set 1, E7335, New England Biolabs). Briefly, eluted RNA was deprived of ribosomal RNA using the NEBNext rRNA depletion kit (BE7405L, New England Biolabs), fragmented before the synthesis of single and double strand cDNA and then adapters were ligated.

For both types of libraries, library size distribution and quality was evaluated on the Agilent 4200 TapeStation System (Agilent Technologies Ltd). Finally, libraries were pooled in equimolar concentration based on Qubit (Invitrogen) quantification, and sequencing was performed on a NextSeq™ 550 (Illumina) with 2×76 cycles, following the manufacturer’s instructions. Quality control of the raw sequencing data was done using FastQC (v0.11.9, Babraham Institute, Babraham, UK). Subsequently, Trimmomatic (v0.39) was employed to eliminate low-quality reads and conduct adapter trimming. The extracted RNA sequences were aligned with the human reference genome (hg19) using STAR (v2.7.0) (https://github.com/alexdobin/STAR/releases). Transcript abundances were quantified with RSEM (v1.3.3) (https://deweylab.github.io/RSEM/) and gene expression levels normalized to transcripts per million (TPM).

### RT-qPCR analysis

The RNA was retro-transcribed into cDNA using the ImProm-IITM (A3800, Promega), according to the vendor’s instruction. The cDNA was then diluted 1:5 with water and 5% of the diluted material was analyzed by RT-qPCR analysis using the POWER SYBR GREEN PCR master mix (4367659, ThermoFisher Scientific). The values obtained for each immunoprecipitated RNAs were normalized over the respective input material and plotted in a histogram, as relative fold enrichment.

The following primers have been used for RT-qPCR analysis:

RPL26 Forward primer 5’ -GGCTAATGGCACAACTGTCCAC- 3’; Reverse primer 5’ - GGCGAGATTTGGCTTTCCGTTC- 3’.
RN7SL1 Forward primer 5’- CATCAATATGGTGACCTCCCG- 3’; Reverse primer 5’- GTTTTGACCTGCTCCGTTTC- 3’.

### Immunofluorescence analysis

Cells were fixed in 4% paraformaldehyde (12606, Cell Signaling Technology) dissolved in PBS for 10 minutes at room temperature, followed by 3 washes with PBS 1x of 5 minutes each. Cells were then permeabilized by adding 0.1% TRITON-PBS for 10 minutes at R.T. and washed 3 times with PBS 1x for 5 minutes each washing step. Aspecific sites were blocked with 2 % BSA-PBS 1x for 30 minutes at R.T., prior adding the anti-FLAG M2 (F1804, Merck Life Science) primary antibody diluted in 2 % BSA-PBS 1x for 2 hours at R.T. After the incubation, 3 washes with PBS 1x of 5 minutes each were performed, followed by the staining with the secondary antibody (Alexa 488 (A11029 and A11034, Invitrogen) or Alexa 555 (A31572, Invitrogen) for 1 hour at R.T.. At the end, 3x washes in PBS of 5 minutes each were performed and the coverslips mounted with antifade mounting medium with DAPI (EN53003M010, Enzo Life Sciences).

### Live-cell imaging

A TET-ON system for the tetracycline-inducible expression of HEK293T cells was generated by transfecting with ViaFECT (E4981, Promega) a mammalian expression plasmid expressing the transactivator rtTA3. Cells stably integrating the plasmid were selected through puromycin selection. These cells were then transiently transfected overnight with the pRetroX GFP-HEPN. The following day, the cultivation media was replaced with fresh medium supplemented with 1 μg/mL doxycycline to activate the TET-ON system. The expression of GFP-HEPN was acquired in live cells at the indicated time points, using the Evident-Olympus IX83 microscope.

### Protein extraction and Western Blot analysis

Proteins were extracted using RIPA Buffer (10 mM Tris-HCl pH 8.0, 1% NP-40, 150 mM NaCl, 0.1% SDS, 0.1% NaDeoxycholate, 1mM EDTA) complemented with cocktail protease inhibitor (11836153001, Merck Life Science). Cells were lysed on ice for 15 minutes at 4 °C and then spun down at 15000 g for 15 minutes. The supernatant containing the protein extracts were transfer into a clean 1.5 mL tube and protein concentration quantified by Bio-Rad Protein Assay Dye (5000006, BIO-RAD Laboratories). For Western Blot analysis, 20 μg of protein whole cell extracts were resolved by SDS-PAGE gel and then transferred into a nitrocellulose membrane. The membrane was blocked with 5% milk dissolved in 0.05% TWEEN-TBS 1x and then incubated over nigh at 4 °C with one of the following primary antibodies: anti-FLAG M2 (F1804, Merck Life Science); Anti-SACSIN antibody (EPR11864, Abcam); anti-GAPDH (14C10, Cell Signalling); anti-Vinculin (A2752, ABclonal). The following day, after three washing steps of 5 minutes with 0.05% TWEEN-TBS 1x, the membrane was incubated for 1 hour with the HRP-conjugated secondary antibody (1706515 and 1706516, BIO-RAD Laboratories), washed for additional three washing steps of 5 minutes with 0.05% TWEEN-TBS 1x and then revealed by chemiluminescence using the clarity western ECL substrate (1705061, BIO-RAD Laboratories) and recorded with the ChemiDocTM Imaging System (12003153, BIO-RAD Laboratories).

### GST-pulldown assay

To preserve the proper RNA-folding^43^, the synthetic biotinylated RNA was boiled for 2 minutes at 95 °C and then put on ice for 5 minutes. Subsequently, the RNA was resuspended in the pulldown reaction buffer (PD-buffer: 2 mM MgCl_2_, 100 U/mL RNAse inhibitor, protease inhibitor, 0.05% BSA, 0.2% NP-40, PBS 1x) and let it stand for 20 minutes at room temperature. For the assay, 10 pmol of RNA was denatured and refolded in a final volume of 550 μL of PD buffer, according to the abovementioned protocol. Afterwards, 10% of the material was taken as input and the remaining 90% divided in two 1.5 mL tubes containing 20 μg of either GST-HEPN or GST-UBI-SR1, previously quantified by Coomassie staining into a final volume of 500 μL. Samples were rocked on a wheel for 1 hour at room temperature and afterwards beads were pulled down by centrifugation and washed 3 times with PD-buffer and the bound RNA extracted with the Miny RNAse extraction kit (R1054, Zymo Research). The RNA was eluted in 30 μL of nuclease free water and 2 μL used for the RNA dot blot assay.

### *cat*RAPID analysis

*cat*RAPID prediction to determine the interaction propensity of SACSIN toward cellular RNA transcripts were conducted using the *cat*RAPID omics v2.1 webfold^28^. The interaction propensities of the amino acidic sequence of canonical full length SACSIN (Q9NZJ4-1), the UBL (SACSIN amino acids 1-84) and HEPN (SACSIN amino acids 4441- 4579) domains were investigated against human protein-coding RNAs. A positive interaction propensity is considered when the Z-score was > 0.5.

### Generation of Venn diagrams

Venn diagrams were generated using the following webtool: Oliveros, J.C. (2007-2015) Venny. An interactive tool for comparing lists with Venn’s diagrams. https://bioinfogp.cnb.csic.es/tools/venny/index.html

### Molecular Modeling

The structure of HEPN dimer in complex with the following RNA sequences were predicted using AlphaFold 3 ^30^. To get a feasible simulated system size, we used RNA sequences of 21 nucleotides.

Sequence of the predicted RNA interactor: 5’ – GGGCCUGGAGCCGGAUCUAA- 3’.
Sequence of the non-predicted RNA interactor: 5’ - GGGGUGGGGGAGGACGCCG - 3’.

The predictions were of good quality for the protein, but scarce for what concerns the RNA structure. However, the predicted structures were sufficiently good to run stable molecular dynamics simulations using Gromacs vers. 2021.3 ^44^. The complexes were solvated with TIP3 water molecules and 150mM of KCl ions were added using CharmmGUI^45^. We used the Charmm36m force field^46^ and the Particle Mesh Ewald to describe the electrostatic interactions. The structures were minimized for 1000 steps, then a 2.5 ns NVT simulation was performed while harmonically restraining the complex to the initial coordinates. Eventually, NPT simulations were performed using a timestep of 2 fs for 500 ns.

### Biolayer interferometry

Biolayer interferometry (BLI) experiments were performed using an Octet Red instrument (ForteBio, Inc., Menlo Park, CA) at 25 °C. Binding assays were carried out in 20 mM potassium phosphate buffer (pH 7.4) containing 150 mM KCl and 0.05% Tween-20. Streptavidin-coated biosensors were loaded with 0.5 µg/ml of biotinylated RNA aptamer and subsequently exposed to increasing concentrations of protein, ranging from 75 nM to 10 µM. Dissociation constants (K_D_) were determined by fitting the steady-state response (wavelength shift upon binding) as a function of protein concentration. Each assay was repeated three times, with measurements performed in triplicate.

### Protein purification

*Escherichia coli* BL21-Gold (DE3) harbouring pGEX4T-HEPN plasmid, coding for the fusion protein GST-HEPN, or PGEX6P1-Ubl-sr1(SIRPT1), coding for GST-UBL-SR1, was grown at 37 °C in LB medium supplemented with 100 µg/mL ampicillin. When OD_600_ reached 0.6, the expression was induced by addition of 0.1 mM isopropyl 1-thio-β-d-galactopyranoside (IPTG) for 4 hours at 30 °C. Upon harvesting, bacterial pellet was washed with PBS (10 mM Na_2_HPO_4_, 1.7 mM KH_2_PO_4_, 136 mM NaCl, 2.6 mM KCl) and resuspended with lysis buffer (PBS supplemented with 1 mg/mL lysozyme, cOmplete™ EDTA-free protease inhibitor mixture (11836170001, Roche), 5 mM MgCl_2_, 10 µg/ml DNase, 2 mM 1,4-dithiothreitol), and disrupted by sonication (Bandelin HD 2070) on ice. The supernatant was collected after centrifugation at 20.000 g, 4 °C for 30 min, filtered using 0.2 μm filter and rocked for 1 hour at room temperature with Glutathione Sepharose 4B, previously equilibrated in PBS. After washing three times with PBS, the resin was stored at 4 °C and used for the GST-pulldown experiments. For the biophysical assays, the GST-tag of HEPN protein was removed by digestion with 1 U/mL thrombin protease (27084601, Cytiva) upon rocking overnight at room temperature. The mixture containing cleaved HEPN protein and thrombin was collected from the resin using PBS as elution buffer. The thrombin protease was removed using HiTrap™ Benzamidine FF column (17514302, Cytiva). Then, HEPN protein was eluted in two steps using PBS first and PBS-1M (10 mM Na_2_HPO_4_, 1.7 mM KH_2_PO_4_, 1 M NaCl, 2.6 mM KCl) later. A further purification was obtained by gel filtration using a Superdex 75 10/300 gl column (17517401, Cytiva) on a FPLC □KTA Pure 25 (Cytiva), eluting with 20 mM phosphate buffer (10 mM Na_2_HPO_4_, 10 mM NaH_2_PO_4_, pH 7.4), 0.15 M NaCl, 5% (w/v) glycerol. The purity of the protein was verified by SDS-PAGE followed by Coomassie staining. Protein was stored at − 80 °C.

### Sample preparation for fluorescence and circular dichroism experiments

A 20 µM RNA solution was denatured at 95 °C in nuclease-free water for 2 minutes and then cooled on ice for 5 minutes. After cooling, it was diluted to obtain a 6 µM RNA sample in 20 mM phosphate buffer, 150 mM NaCl, 5% glycerol, RNase inhibitor (40 U/µL), and 2 mM MgCl□.

After 20 minutes, the refolded RNA was mixed with the HEPN samples at the required concentrations to achieve the desired composition of both components, maintaining RNA-to-protein volume ratio of 1:2.

### Fluorescence Spectroscopy

Fluorescence experiments were performed by using a JASCO FP6500 spectrofluorometer (JASCO Corporation, Tokyo, Japan) at T=20 °C. Intrinsic fluorescence experiments were performed registering emission spectra after excitation at 295 nm (excitation slit width = 3 nm, emission slit width = 3 nm, response 2 s, scanning rate 100 nm/min) using a quartz cell with a path length of 1 cm.

Thioflavin-T (ThT) fluorescence (12 µM ThT, 20 °C; λ_exc = 450 nm, λ_em = 485 nm) was measured for ThT alone and after incubation with HEPN samples obtained at the start and at the end of the condensate-formation kinetics.

### Circular dichroism

CD spectroscopic measurements were made by using a JASCO J-810 spectrometer equipped with a temperature control unit. A quartz cell with a path length of 0.5 mm was used for recording at 20 °C the spectra (200-320 nm) of a sample containing 2 µM RNA in the absence or in the presence of 3 µM, 5 µM, 9 µM or 20 µM HEPN. Spectra were acquired with a 20 nm/min scan rate, band width 2 nm, response 16 s and data pitch 0.5 nm. CD spectra of the protein alone were also recorded in the same conditions for each concentration used. Samples were corrected using the respective blank.

### Multi Angle Light Scattering (MALS)

Protein solutions were prepared at 23 μM in 10mM phosphate buffer, 150 mM NaCl, and 5% (v/v) glycerol. Where indicated, RNA was added to the protein solution at 0.40 μg/μL (low RNA) or 0.44 μg/μL (high RNA). For measurements, 170 μL of the protein solution (with or without RNA) was mixed 1:1 with 170 μL of PEG6000 (30% w/w; Farmalabor) prepared in the same buffer, yielding a final volume of 340 μL. The final concentration upon mixing were: HEPN domain 11.5 μL, PEG6000 15% w/w, and RNA 0/0.20/0.22 μg/μL for the no-RNA, low-RNA, and high-RNA conditions, respectively. All solutions were handled under dust-free conditions and loaded into cylindrical quartz cells. Multi-angle light scattering measurements were performed on a Brookhaven Instruments BI-200SM goniometer equipped with a 50□mW He–Ne laser (λ□=□632.8□nm) and a Brookhaven BI-9000 correlator. Cells were mounted in a thermostated compartment maintained at 20.0□°C (± 0.1□°C) via a recirculating bath.

Kinetic acquisition: Time-lapse scattering at □=30° was collected every 3□min for 2□days to monitor condensate formation kinetics.

Angular acquisition: For absolute form-factor analysis, the scattering intensity I(q) was measured at 27 angles, corresponding to q spanning 5.3–25.2□µm□¹. The magnitude of the scattering vector was defined as q = 4πn/λ sin(□/2), with n the solvent refractive index At □=30°, q≈6.8μm^ (−1).

Measured intensities were converted to the absolute Rayleigh ratio R(q) using toluene as the standard at □=90°, with R_tol (90°, λ = 632.8nm) = 1.4 10^(−5) 〖cm〗10^(−1). The sample Rayleigh ratio at angle □ was obtained from R(q(θ)) = (I(q(θ)) sinθ/ (I_tol (90°)) R_tol (90°), where I_tol (90°) is the toluene intensity measured under identical optical geometry. For dilute solutions, the angular dependence of the absolute intensity was analyzed using R(q) = KCMP(q), where K is the optical constant, C the solute concentration, M the intensity-weighted average molar mass of the scatterers, and P(q) the form factor. Spherical form factors were used to extract the gyration radius R_g and infer particle morphology. For HEPN alone in 15% PEG6000 data were best described by a spherical specie with R_g 10^ ((1)) = 440nm and R_g 10^ ((2)) = 2.6μm, while the two samples with RNA were analysed by using a spherical specie with R_g (low RNA) = 330nm and R_g (high RNA) = 250nm, respectively. Best-fit form factors for the three conditions are shown as continuous lines in Figure□4A, together with the corresponding absolute-scale MALS data. The kinetics trace at □=30° (absolute intensity versus time) is reported alongside the form-factor at 48h for each sample.

## DATA AVAILABILITY STATEMENT

RNA-Seq data were deposited in the GEO public repository database with the following number: GSE312176.

## ACKNOWLEDGMENTS AND FUNDINGS

We thank E.A. Calcagno for technical support and Dr. V. Stagni for carefully reading the manuscript.

This work was supported by Fondazione Telethon and Associazione ARSACS OdV (Fall Seed Grant 2023 #GSA23L002)”. The research conducted in the laboratory of RG was also supported by Unione europea- Next Generation EU, Missione 4 Componente 1, CUP B53D23016490001.

## AUTHOR CONTRIBUTIONS

R.G. conceptualize the paper, designed and contributed to the execution of the experiments, analysed the data, wrote the manuscript and provided fundings; E.Z. conceptualize and carried out the BLI experiment and wrote the manuscript; I.M., A.C. and G.C. performed, collected and interpretated the cellular and molecular biology experiments. L.L., R.C., M.R.M, M.A.C., R.P., S.V. and V.M. conceptualize, performed, collected and interpretated the biophysical experiments. V. M. designed and carried out the experiment of molecular dynamics and the AlphaFold 3 predictions. R.V., N.C. and A.G. performed and analysed the RNA-Seq data.

All authors actively contributed to this work, revised the manuscript and agreed to its content.

## Supplementary figure and file legends

**Supplementary Figure 1.**
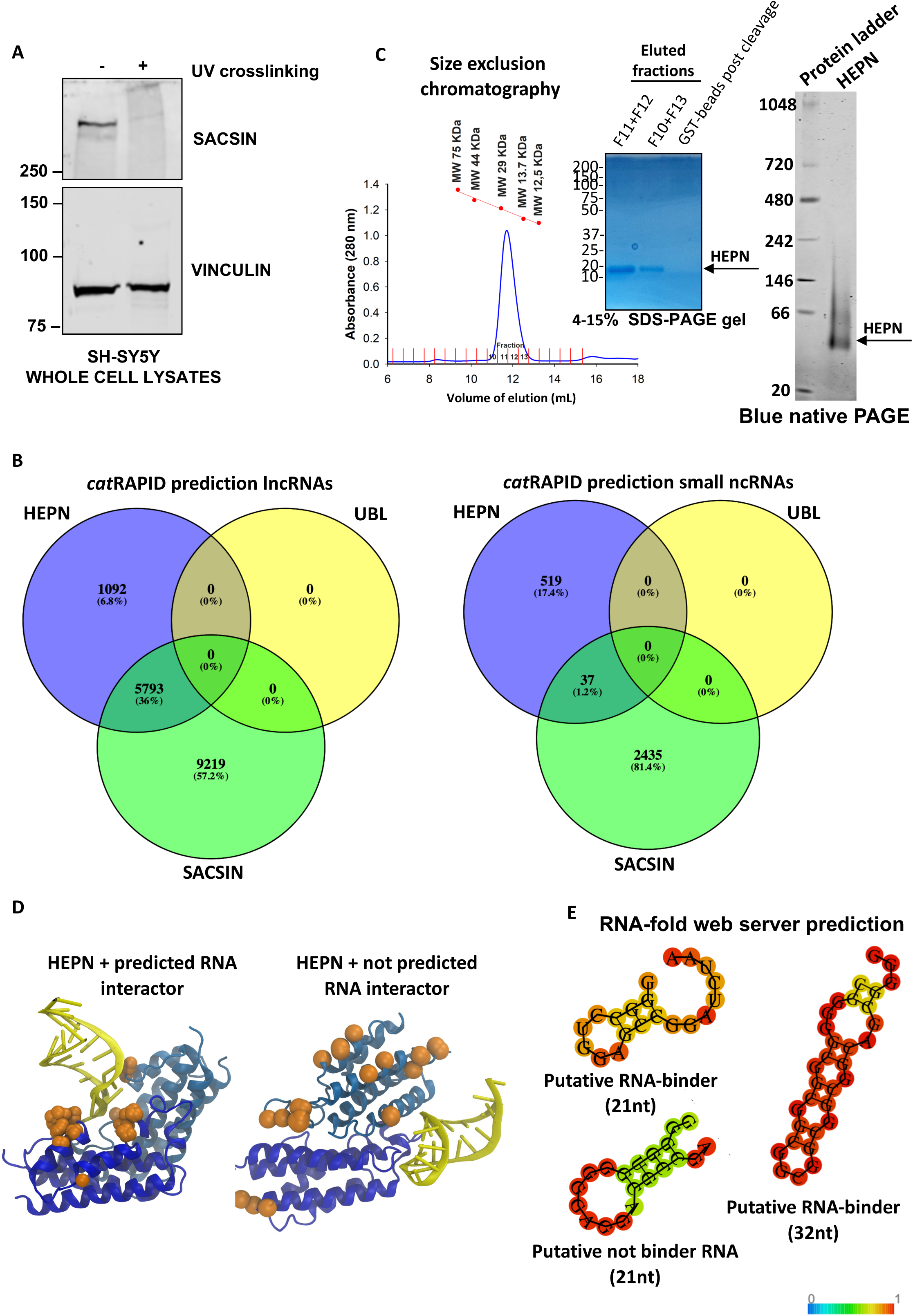
**A)** Western blot analysis of SH-SY5Y whole cell lysates in normal condition and upon UV-crosslinking. In response to UV-crosslinking, the detection of SACSIN is strongly reduced and it is possible to observe also a weak band at a very high molecular weight band detected by SACSIN antibody. The displayed figure is the representative image of one of the three performed biological replicates (n = 3). **B)** *cat*RAPID predictions of SACSIN and SACSIN domains with either long non-coding RNA (lncRNA) (left) or small non-coding RNAs (ncRNAs) (right). **C)** Left. Size exclusion chromatography (SEC) profile of the HEPN domain (Molecular weight = 15 kDa) after the thrombin cleavage of the GST-tag (blue line). The molecular weight calibration curve (red) obtained with the protein gel filtration calibration kit (Cytiva) indicates that HEPN elutes in its dimeric form. The purity of the purified HEPN was confirmed by Coomassie staining on an SDS-PAGE gel. Right. Native Blue PAGE analysis of HEPN domain confirmed that HEPN is a natural dimer. **D)** Snapshots from molecular dynamics trajectories of dimeric HEPN in the presence of a specifically interacting RNA (left) and not specifically interacting RNA (right). The Cα of the amino acids involved in the interaction with the RNA during the respective trajectories are displayed as orange spheres. **E)** Predicted secondary structure of the RNA sequences used for the molecular dynamics and the GST-pulldown assay.

**Supplementary figure 2.**
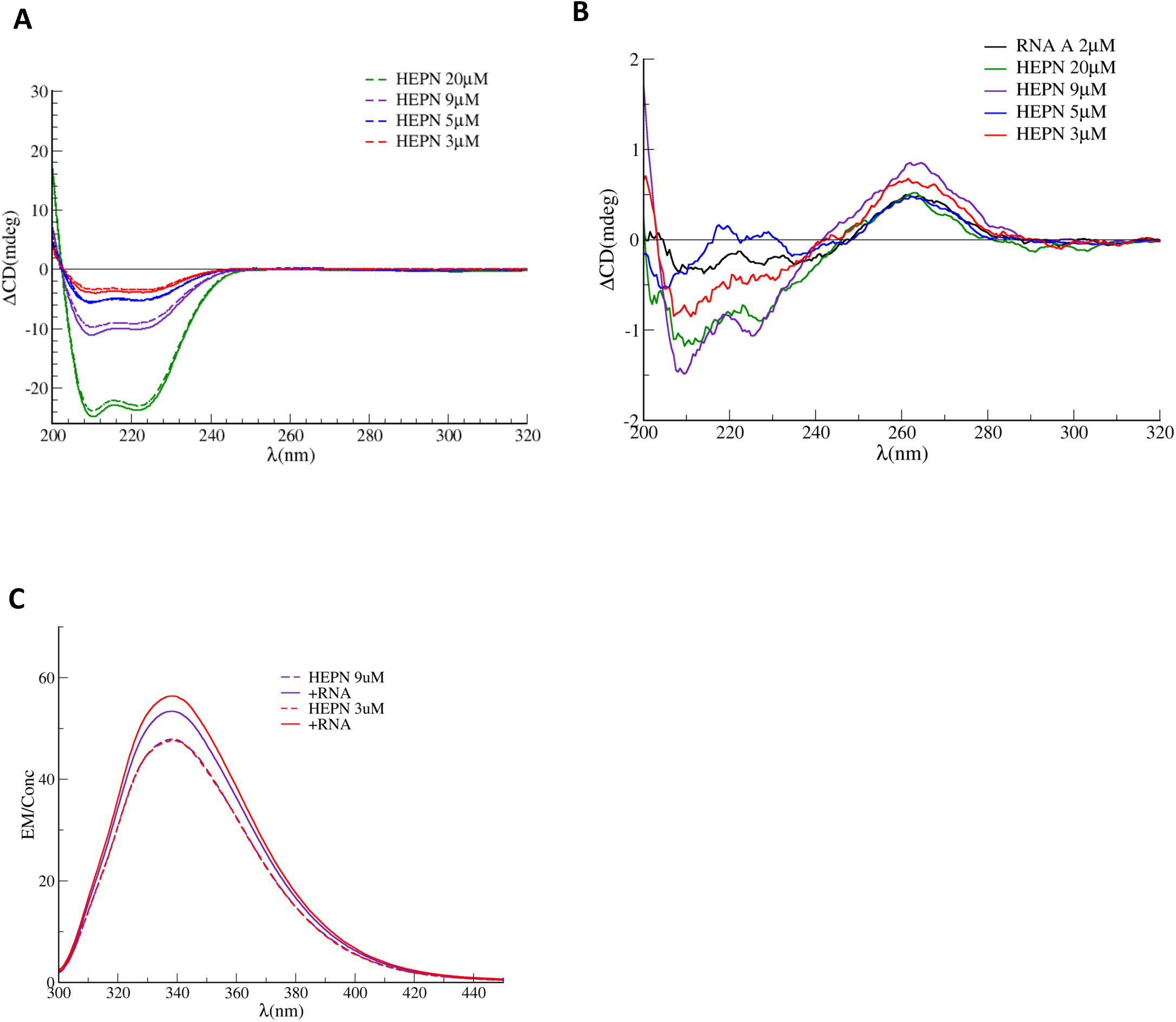
**A)** circular dichroism (CD) experiments of HEPN with and without RNA. Comparison of Far-UV and Near-UV CD spectra of HEPN obtained by subtracting the RNA signal from the samples containing HEPN at varying concentrations alone (continuous lines) and with a fixed RNA concentration of 2 μM (dashed lines). The shape of the CD spectra did not change in the presence of the RNA, suggesting that there is no alteration of the HEPN secondary structure. **B)** Same experiment described in A but with the focus on RNA CD. Difference in the CD spectra obtained by subtracting the signal of the protein alone from that of the HEPN+RNA samples at various protein concentrations: 3 μM (red), 5 μM (blue), 9 μM (violet), and 20 μM (green). The CD spectrum of RNA alone is shown for comparison (black). The results showed fluctuations (“flickering”) in the difference signal across protein concentration and the absence of a proportional increase in the difference signal, both in the Far- and Near-UV regions. **C)** Emission fluorescence spectra of HEPN after excitation at 295 nm that showed very mild differences in the intrinsic fluorescence of the protein in the presence of the RNA oligo. No clear shift of the curve was observed when the HEPN and the RNA were mixed in the same reaction tube compared to the HEPN alone. In the presence of the RNA, only a marginal increase of the intrinsic fluorescence of HEPN can be measured.

**Supplementary Figure 3.**
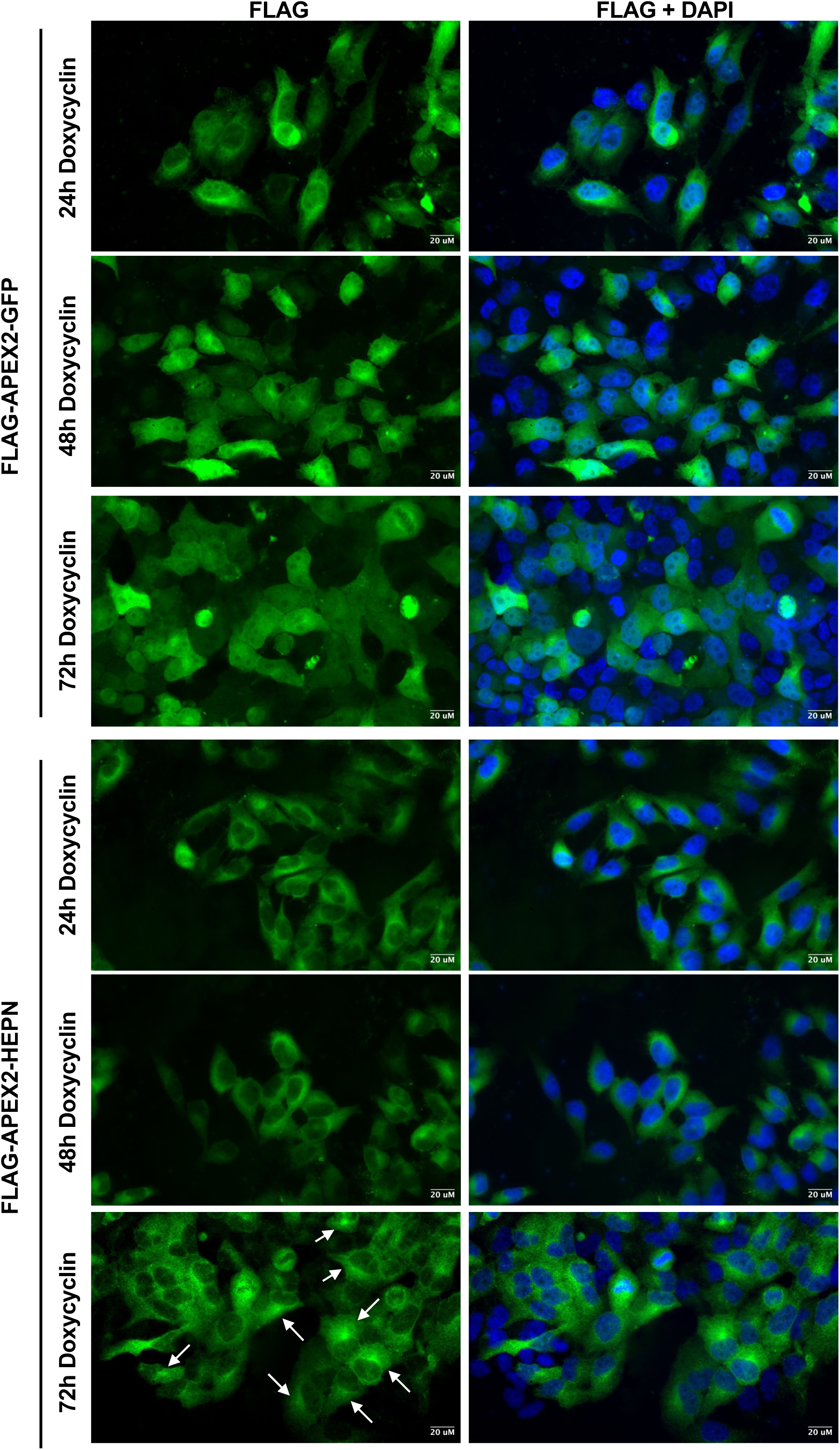
Immunofluorescence analysis of Hela Flp-In^TM^ T-Rex cells expressing either FLAG-APEX2-GFP or FLAG-APEX2-HEPN, upon doxycycline administration over time. Cells were induced for the indicated time points and then fixed with 4% PFA followed by immunofluorescence staining with a-FLAG antibody and DAPI. The white arrows indicated the cells where we observed a localization of the protein in the ER-proximity. The displayed figures are representative images of one of the two performed biological experiments (n = 2).

**Supplementary Figure 4.**
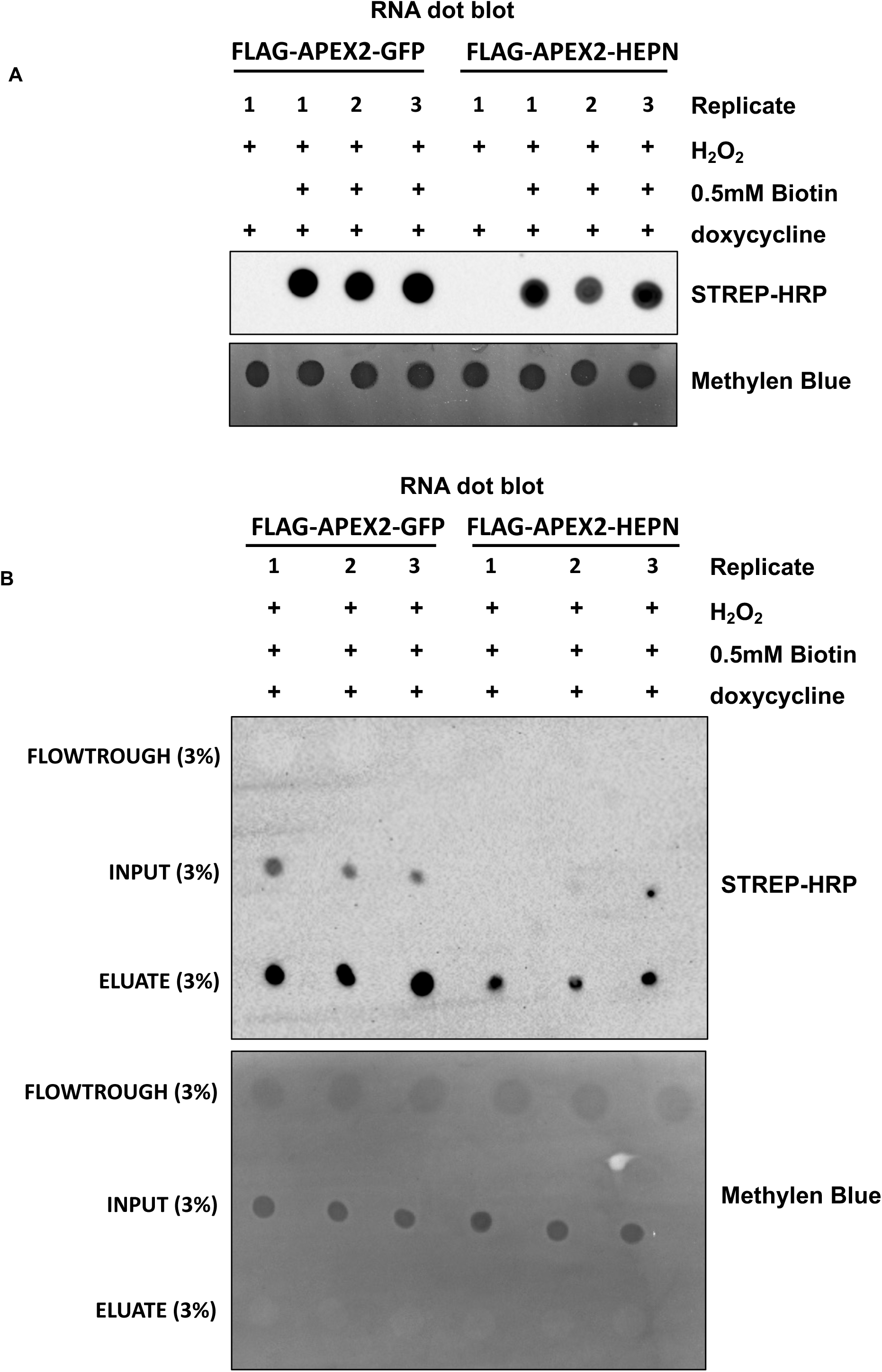
**A)** RNA dot blot analysis of the biotinylated RNAs recovered from three independent biological replicates performed upon 72h of 1μg/mL doxycycline administration in Hela Flp-In^TM^ T-REx FLAG-APEX2-GFP and FLAG-APEX2-HEPN cells. **B)** Streptavidin pulldown to affinity purified the biotinylated RNAs that have been subsequently analyzed by next generation sequencing.

**Supplementary Figure 5.**
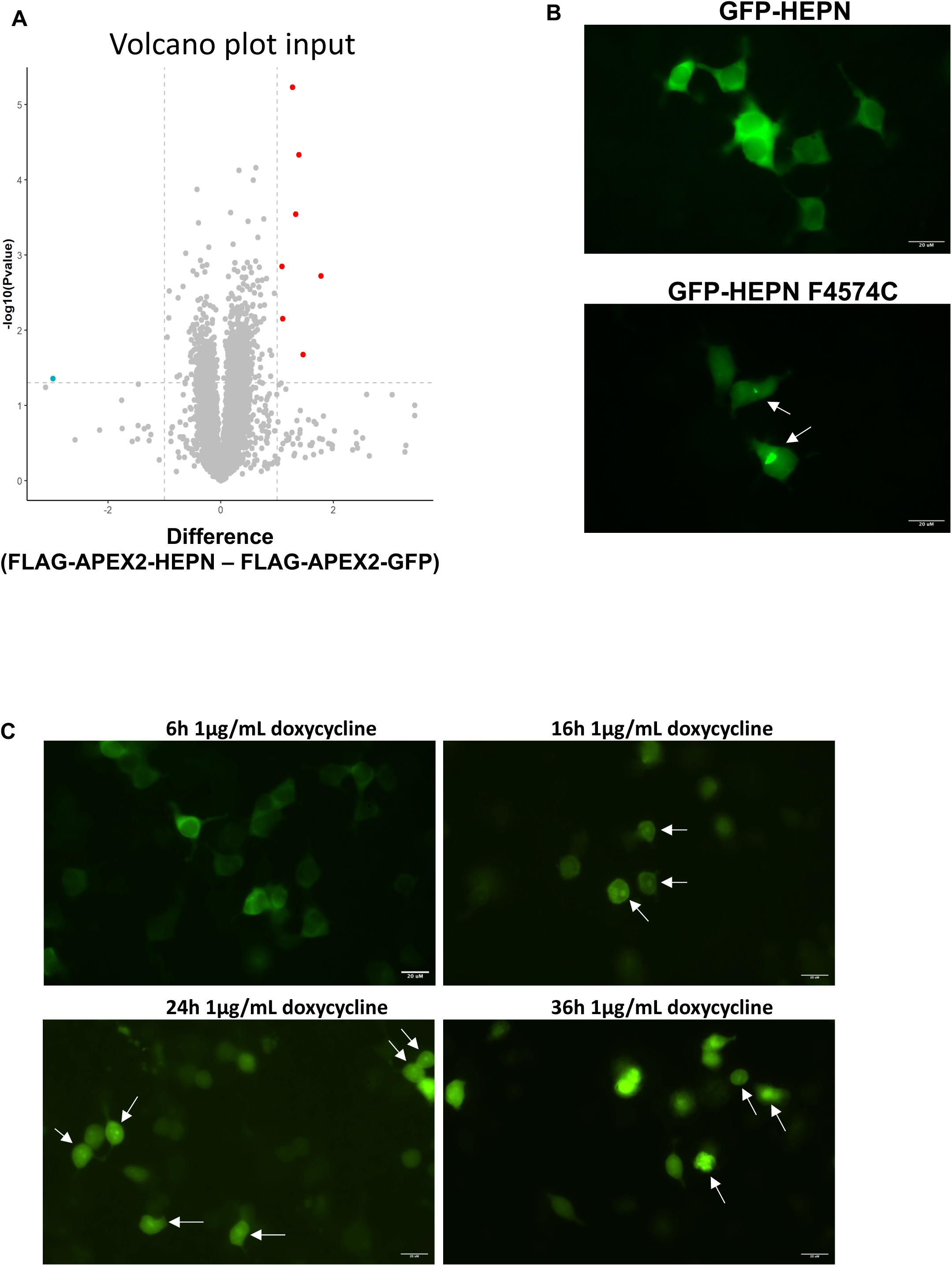
**A)** Volcano plot analysis of input material derived from FLAG-APEX2-HEPN and FLAG-APEX2-GFP cells. RNA transcripts significantly enriched (log_2_ Fold change (Fc) > 1; two-sided T-Test Pvalue < 0.05) in FLAG-APEX2-HEPN and FLAG-APEX2-GFP cells are coloured in red and blue, respectively. **B)** Live imaging of HEK293T cells co-transfected with either GFP-HEPN, the mutated F4574C or N4549D forms and a rTTA3-containing plasmid. Cells were induced with 1 μg/mL doxycycline to promote the transcription of GFP-HEPN mRNAs and images were taken after 6h of doxycycline administration. The white arrows indicated cells where we observed protein condensates. The displayed figures are the representative images of one of the three performed biological replicates (n = 3). **C)** Live imaging of HEK293T cells carrying a tetracycline-inducible plasmid for the expression of the transactivator rtTA3 that were transiently transfected with GFP-HEPN. Cells were induced with 1 μg/mL doxycycline to activate the TET-ON system, promoting the transcription of GFP-HEPN mRNA. Images were taken at the indicated time points. The white arrows indicated cells where we observed protein condensates. The displayed figures are representative images of one of the two performed biological replicates (n = 2).

**Supplementary Figure 6.**
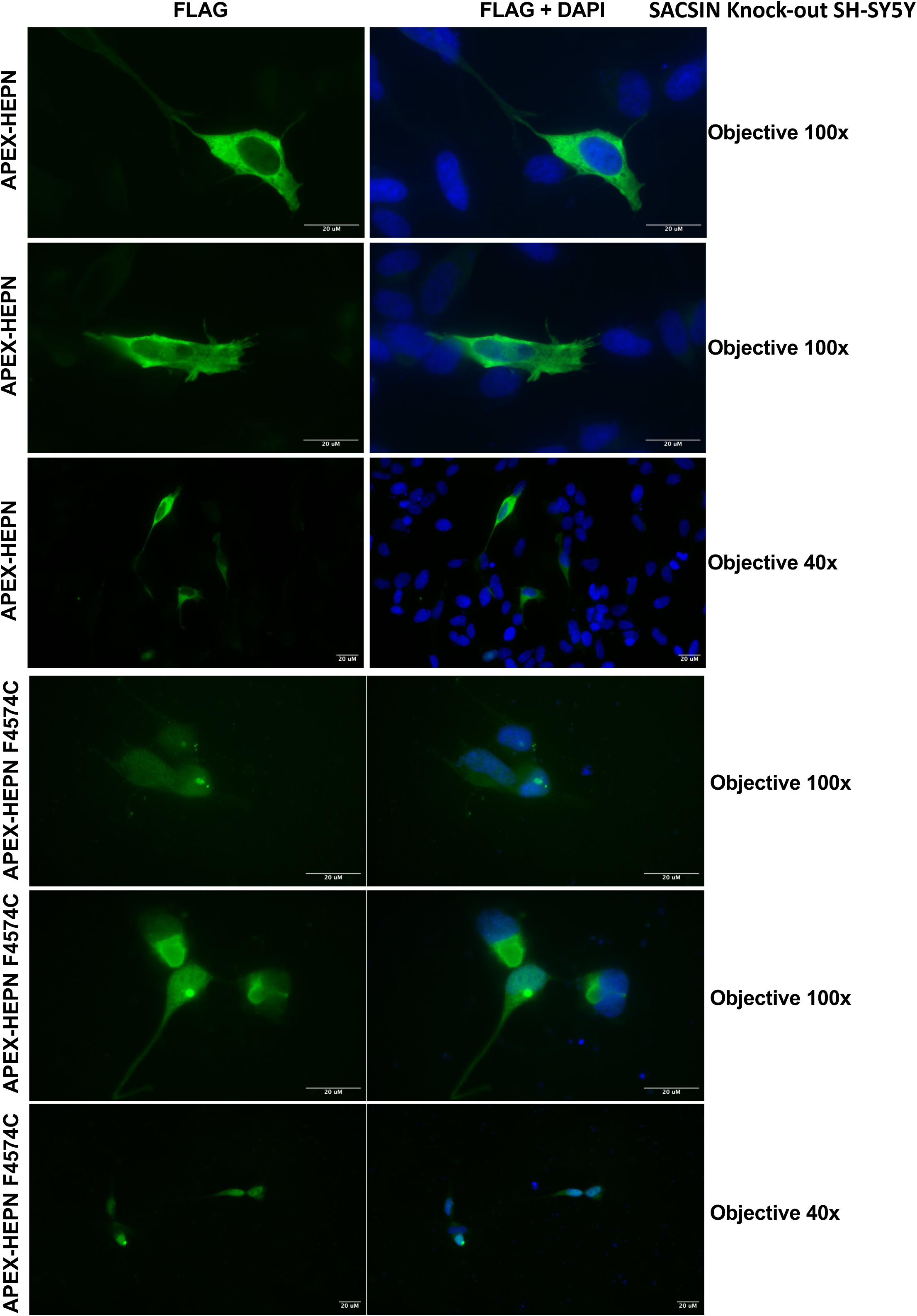
Additional immunofluorescence images of SACSIN knock-out SH-SY5Y cells transfected with either FLAG-APEX2-HEPN or the mutated F4574C form, as described in Figure 4D. Images were taken at both 40x and 100x magnification.

**Supplementary Figure 7.**
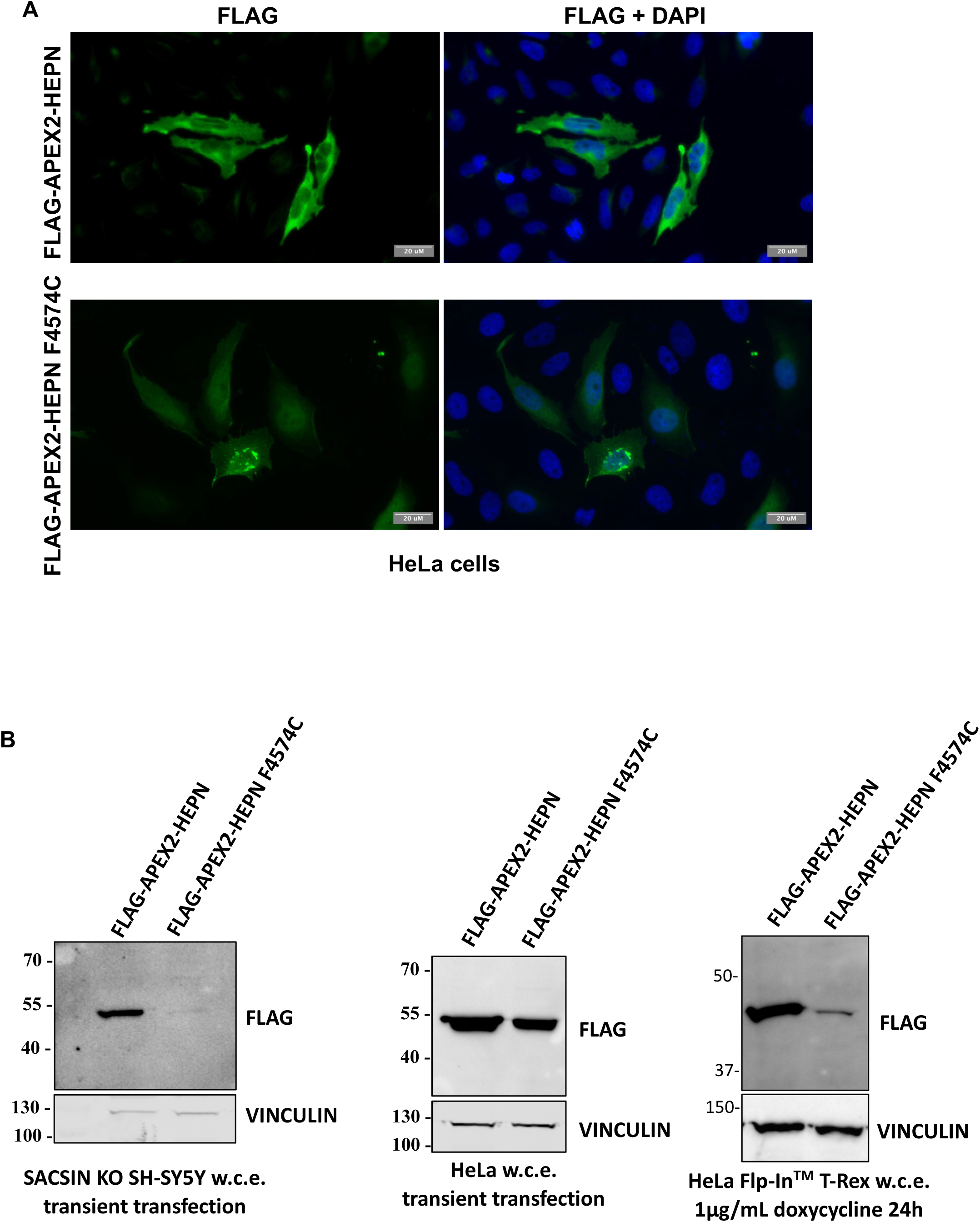
**A)** HeLa cells transfected with either FLAG-APEX2-HEPN or the mutated F4574C form. Transfected cells were fixed and stained with anti-FLAG and DAPI to assess the cellular localization of FLAG-APEX2-tagged proteins. FLAG-APEX2-HEPN was expressed in the cytoplasm while the F4574C showed a nuclear localization and the presence of protein condensates. The displayed figures are representative images of one of the three performed biological replicates (n = 3). **B)** Western blot analysis of SACSIN knock out (KO) SH-SY5Y (left) and HeLa (middle) cells transiently transfected with a plasmid for the expression of the FLAG-APEX2-HEPN protein. (Right) Western blot analysis of HeLa Flp-In^TM^ T-REx expressing either FLAG-APEX2-HEPN or the mutant FLAG-APEX2-HEPN F4574C after 24h of 1 μg/mL doxycycline administration. The displayed figures are the representative images of one of the two performed biological replicates (n = 2).

**Supplementary File 1. Prediction of an RNA sequence bound by SACSIN HEPN.** Sequence alignment analysis using Clustal Omega^29^ of the strongest RNA interactors predicted by *cat*RAPID (Z-score > 3.5) and in common between SACSIN and SACSIN HEPN domain.

The sequence of RNA transcripts with a Z-score < 0 were used as control references.

The sequences reported in the document are the HEPN-interacting RNA regions predicted by *cat*RAPID.

**Supplementary File 2. RNA-Seq data of input and eluted materials of APEX-Seq.** Differentially regulated genes are expressed as follow: FLAG-APEX2-HEPN (UP) and FLAG-APEX2-GFP (DOWN) cells. Data were quantified in Transcript per million (TPM). TPM low indicates if the TPM value was < 10. Abbreviations present in the file: n. replicates GFP = number of replicates in which the RNA was found; sd = standard deviation; FC = Fold Change.

Significant RNAs were considered if they had a log_2_ fold change (FC) FLAG-APEX2-HEPN – FLAG-APEX2-GFP > 1 or < 1 and a two-sided T-Test Pvalue < 0.05. APEX-Seq data of input and eluate materials are reported in the datasheets “Input” and “Eluates”, respectively. Datasheet “HEPN significant RNA interactors” displays only the significant RNA transcripts found associated with FLAG-APEX2-HEPN domain. Datasheet “GO eluate UP” and GO eluate DOWN” include the gene ontology enrichment analysis of the significant RNA transcripts associated with FLAG-APEX2-HEPN and FLAG-APEX2-GFP, respectively.

**Graphical abstract** created in BioRender. Vazzana, R. (2025) https://BioRender.com/rjn557o

